# Transformer-based Deep Learning for Glycan Structure Inference from Tandem Mass Spectrometry

**DOI:** 10.1101/2025.07.02.662857

**Authors:** Ejas Althaf Abtheen, Arun Singh, Shyam Sriram, Changyou Chen, Sriram Neelamegham, Rudiyanto Gunawan

**Affiliations:** Department of Chemical and Biological Engineering, University at Buffalo-SUNY, Buffalo, NY 14260; Department of Bioengineering and Biomedical Engineering, University of Pennsylvania, Philadelphia, PA 19104; Department of Computer Science, University at Buffalo-SUNY, Buffalo, NY 14260; Cell, Gene, and Tissue Engineering Center, University at Buffalo-SUNY, Buffalo, NY 14260

## Abstract

Glycans play critical roles in diverse biological processes, but their structural analysis by tandem mass spectrometry (MS/MS) remains a major challenge due to their branched structure and stereochemistry. Traditional computational methods, such as database searching, are constrained by the scope of existing libraries and can be computationally intensive. While recent deep learning models have advanced the field, they often struggle to capture the complex, long-range dependencies within MS/MS spectra required for accurate inference. To address these challenges, we present GlycoBERT and GlycoBART, novel transformer-based models for glycan structure prediction from MS/MS data. GlycoBERT, a sequence classifier, achieves 95.1% structural accuracy on test data, surpassing the current state-of-the-art deep learning model, CandyCrunch. However, classification-based methods are inherently limited to predicting structures present in the training data. To overcome this, we developed GlycoBART, a generative sequence-to-sequence model capable of *de novo* glycan inference. On independent validation datasets, both models demonstrate robust performance. Critically, when applied to an MS/MS dataset from human embryonic kidney cells, GlycoBART generated two *de novo* glycan structures absent from the training set, one of which is a novel structure not catalogued in major glycan databases. Together, GlycoBERT and GlycoBART establish a new benchmark for glycan analysis, offering a powerful framework that enables more accurate and comprehensive exploration of glycan diversity and discovery.

## INTRODUCTION

Glycans, complex carbohydrates covalently attached to proteins and lipids, play a pivotal role in a range of biological processes, including cell-cell communication [1, 2], immune response modulation [3, 4], protein folding [5-7], and pathogen recognition [8]. The structural diversity of these complex carbohydrates, arising from differences in monosaccharide composition, linkage types, branching patterns, and terminal residues, underpins their functional specificity. Aberrations in glycan structures are hallmarks of various diseases, including cancer [9, 10], autoimmune disorders, and infectious diseases [11], underscoring their diagnostic and therapeutic relevance. Thus, systematic characterization of glycan structures or glycomics is essential for advancing our understanding of their functions and harnessing their potential in medicine and biotechnology.

Mass spectrometry (MS) is a highly sensitive and versatile analytical technique widely employed in glycomics for the structural characterization of glycans [12-15]. A common approach involves coupling liquid chromatography coupled with tandem mass spectrometry (LC-MS/MS), which enables the separation of glycan isomers and provides detailed structural information [16-20]. In tandem MS analysis, precursor glycans are ionized and fragmented, generating mass spectra that record the mass-to-charge (m/z) ratios of the resulting fragment ions (see Fig. 1A). These fragmentation patterns typically involve glycosidic bond and cross-ring cleavages, yielding diagnostic ions that reveal key information about the glycan’s monosaccharide composition, branching, linkage types, and terminal modifications [21-24]. The structural features of glycans are then determined by analyzing the m/z values and relative intensities of these fragment ions.

**Fig. 1.**
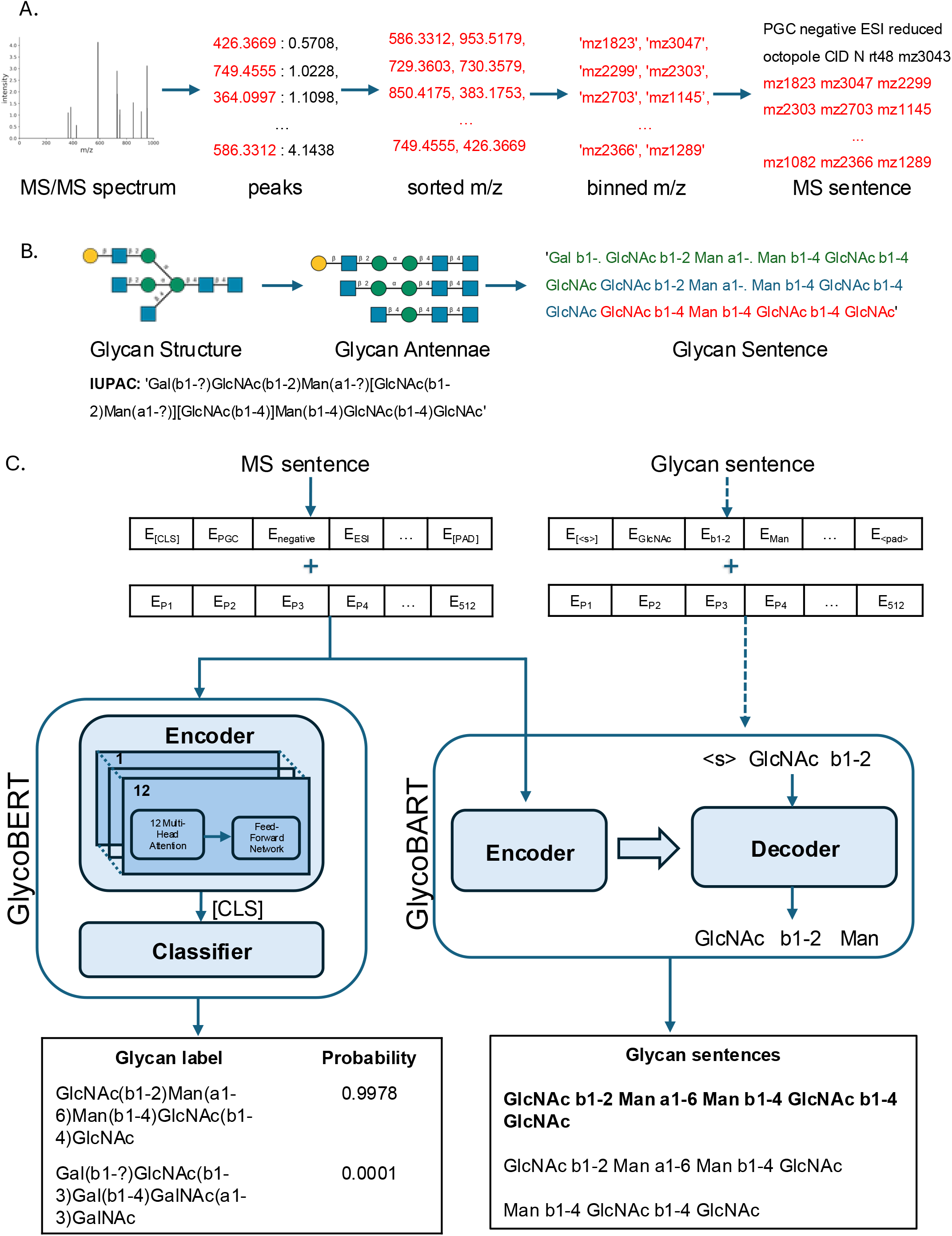
Mass spectra and glycan representation and model architectures for GlycoBERT and GlycoBART. (A) MS sentence representation. An MS/MS spectrum is transformed into an “MS sentence.” This sequence of discrete words encodes experimental parameters, retention time, precursor mass, and spectral data. To incorporate intensity information, fragment ion peaks are ordered by descending intensity, allowing their relative importance to be captured by the model’s positional embeddings. (B) Glycan sentence representation. A glycan structure is linearized into a “Glycan sentence.” Each glycan is composed of its constituent antennae, with each antenna described as a sequence of alternating monosaccharide and linkage words, from the terminal residue to the core. (C) Model training and inference workflows. Both DL models take a tokenized MS sentence as input to a transformer encoder. GlycoBERT, a classification model, passes the encoder output to a classification head that give confidence scores to each of 3590 glycan structures in training dataset. GlycoBART, a conditional generation model, uses a decoder to generate a Glycan sentence. During training (dotted arrows), the decoder is given the correct Glycan sentence as input to learn the mapping. During inference (solid arrows), the Glycan sentence is generated autoregressively, conditioned on the input MS sentence.

However, the inherent complexity of glycan structures, including the presence of isomeric forms and diverse branching patterns, poses significant challenges for manual interpretation of MS data. This complexity necessitates the use of advanced computational tools and specialized glycoinformatics databases to facilitate accurate and high-throughput identification and structural annotation of glycans. A multitude of methods have been developed to predict glycan structures or topologies from tandem mass spectrometry (MS/MS) data, each with distinct advantages and disadvantages [25-32].

One of the earliest and most widely used methods is database search, which involves matching experimental MS/MS spectra to theoretical fragmentation patterns derived from reference glycan structures [25-27]. While effective, this method is inherently constrained by the completeness and accuracy of the database and is unable to predict novel glycan structures that are absent from the reference set. To overcome such limitations, *de novo* sequencing methods have been developed, enabling the reconstruction of glycan structures directly from MS/MS data without reliance on a pre-existing database [28-31]. These approaches generate candidate glycan structures based on precursor mass and fragment ion data, scoring them based on fragment counts, biosynthetic rules, or machine learning-based approaches. Although powerful, these methods are computationally intensive and their accuracy is often constrained by the fidelity of the scoring schemes used to evaluate candidate structures.

More recently, neural network-based models have emerged as a transformative tool for glycan structure prediction from tandem mass spectra. For example, the recent CandyCrunch model employs Convolutional Neural Network (CNN) with dilations and residual connections to classify spectra into glycan classes, followed by post-processing filters to refine structural assignments [32]. However, even with advanced features such as dilated convolutions, CNN-based models may struggle to capture complex, non-local relationships and dependencies within spectral data—factors that are often critical for distinguishing between structurally similar glycans.

To address this challenge, transformer architectures [33] have recently been explored in MS data analysis, particularly in proteomics and metabolomics [34, 35]. Transformers employ self-attention mechanisms to effectively model complex, long-range dependencies within data. These attributes make them particularly well-suited for analyzing mass spectra, where the structural information encoded in the fragmentation patterns often involves relationship between peaks that are non-adjacent.

Building on this transformative potential, we developed GlycoBERT and GlycoBART, two novel transformers-based models based on the BERT (Bidirectional Encoder Representation from Transformers) [36] and BART (Bidirectional and Auto-Regressive Transformers) [37] architectures, respectively. These models represent a significant advance in the application of machine learning to glycomics. GlycoBERT, trained as a sequence classifier, excels in classifying mass spectra into glycan structures with high accuracy. Recognizing the limitations of classification-based methods, we further developed GlycoBART, a generative model capable of de novo glycan structure inference, enabling the prediction of novel glycans beyond the scope of existing databases. GlycoBERT and GlycoBART demonstrated significant improvements in accuracy and robustness compared to the CNN-based model CandyCrunch, particularly in terms of structural inference accuracy. Notably, in an application to an unseen MS/MS data from the analysis of human embryonic kidney cells, GlycoBART predicted two de novo glycans that were absent from the the training dataset, highlighting its potential for novel glycan discovery. Overall, GlycoBERT and GlycoBART set a new benchmark for glycan structure prediction and discovery using tandem mass spectrometry analysis.

## METHODS

### Glycomics MS/MS Dataset

Tandem MS dataset for training GlycoBERT and GlycoBART was adapted from CandyCrunch (version 2, downloaded from Zenodo on April 19, 2024) [32]. The CandyCrunch dataset consists of 510,160 MS/MS spectra—mass-over-charge ratio (m/z) and peak intensities of fragment ions, each annotated with one of 3590 glycan structures.Also included is the m/z of the precursor ion and retention time (RT). To curate the training dataset, we removed all spectra that lacked glycan annotation. We also compiled the experimental metadata, comprising liquid chromatography column type (LC type), glycan type, glycan modification, ion trap, ion mode, fragmentation method, and ionization method, from CandyCrunch (Supplementary Table 10), GlycoPOST [38], and the original publications. MS/MS spectra from Kouka *et al*. study [39] were excluded, allowing their use as an independent validation dataset. The training dataset consisted of 503,446 MS/MS spectra.

### Conversion of MS/MS to MS Sentences

**Fig. 1A** illustrates the conversion of an MS/MS spectrum into a word sequence, termed as “MS sentence”, for use with our transformer models. Each MS sentence encodes key information from a single MS/MS spectrum, including its experimental metadata, normalized retention time (RT), precursor ion m/z, and the m/z values and relative intensities of its fragment ions.

An MS sentence begins with seven words representing experimental metadata, arranged in the following order (with possible words in parentheses): LC type (PGC, C18, HILIC, MGC, other_lc), ion mode (negative, positive, other_mode), ionization method (ESI, MALDI, other_ion), glycan modification (reduced, 2AA, permethylated, PA, native, rapifluor, other_mod), ion trap (linear, orbitrap, amazon, MSD, TOF, octopole, other_trap), fragmentation method (CID, HCD, other_frag), and glycan type (N, O, lipids, free, other_type).

Following the experimental metadata, the next word represents the normalized RT. Here, RT values are scaled by division by the largest RT in the same MS/MS run or by 30, whichever is greater. The resulting normalized RT, ranging from 0 to 1, is then discretized into bins with a width 0.01 and is written as a word starting with ‘rt’ followed by the bin index. For example, a normalized RT of 0.47, which falls in the 48^th^ bin, is written as the word ‘rt48’.

The final set of elements in the MS sentence represent the m/z values, starting with the precursor ion and followed by the fragment ions from the MS/MS spectrum. All m/z values are binned using a width of 0.3 Da over a range of 39.714 Da to 3000 Da—the same binning parameters used in CandyCrunch [32]. Each m/z value is then written as a word starting with ‘mz’ followed by the m/z bin index. For example, an m/z value of 500 is written as ‘mz1535’. As illustrated in Fig. 1A, the fragment ion peaks are sorted in descending intensity. This ordering implicitly encodes the relative peak intensity information into the positions of the m/z words. To filter out noise, only fragment peaks with a relative intensity greater than 0.1% of the total intensity of all peaks in the spectrum are included. Notably, this representation of the fragment ion peaks differs from the vector representation in CandyCrunch. While CandyCrunch representation aggregates multiple m/z peaks that fall into the same m/z bin, our approach treats each peak as a distinct word. Therefore, peaks with m/z values in the same bin will appear as separate words in the MS sentence, their positions differentiated by their relative intensities.

### Processing of Glycan Sentences (GlycoBART)

**Fig. 1B** illustrates the conversion of a glycan structure’s IUPAC representation into a word sequence termed a “Glycan sentence”, designed for training our generative model GlycoBART. In this sentence, a glycan structure is deconstructed into its constituent antennas. Here, each antenna is written as a sequence of words, starting from the terminal monosachharide and proceeding inwards, with monosaccharides alternating with linkages. These individual antenna sequences are then concatenated, separated by spaces, to form the complete Glycan sentence. Consequently, two consecutive monosaccharide words with the sentence mark the boundary between the end of one antenna and the beginning of another. Any unknown linkage information, often marked by a question mark, is denoted by the period symbol (‘.’; see Fig. 1B).

### Tokenization

A custom tokenizer (MSTokenizer.py) was developed for converting MS and Glycan sentences into their corresponding token sequences. The tokenizer utilizes a Word-level approach, mapping each word in a sentence to a unique token ID. Distinct vocabularies were curated for the GlycoBERT and GlycoBART based on this principle.

The GlycoBERT vocabulary is tailored for MS sentences. The vocabulary was curated by aggregating all possible words for the experimental metadata, normalized RT, and m/z values. Additionally, five special tokens required by the BERT architecture ([CLS], [SEP], [UNK], [MASK], and [PAD]) [36] were included. In total, the GlycoBERT vocabulary includes 10,010 unique tokens, comprising 9,870 for m/z, 102 for normalized RT, 33 for metadata, and 5 for special tokens.

The GlycoBART vocabulary builds upon GlycoBERT’s by adding the set of possible words from Glycan sentences. In place of BERT’s special tokens are seven special tokens required by the BART architecture (<pad>, <s>, <\s>,<sep>, <cls>, <unk>, and <mask>) [37]. This results in a total vocabulary of 10,080 unique tokens, comprising 9,870 for m/z, 102 words for normalized RT, 33 for metadata, 68 for glycans, and 7 for special tokens.

### Model Architecture, Training, and Inference

Fig. 1C illustrates the architecture of GlycoBERT and GlycoBART, our transformer-based models for glycan structure classification and generation from MS/MS spectral data, respectively.

GlycoBERT is based on the BertForSequenceClassification from the BERT model [36, 38]. Its core is a BERT encoder with 12 transformer layers, each with 12 attention heads. Following standard BERT practice, a special [CLS] token is prepended to each input MS sentence. The encoder processes the tokenized sequence into 768-dimensional contextual representations. The final representation of the [CLS] token is then used by a classification head which computes confidence scores for each of the 3,590 possible glycan structure classes. During inference, the model outputs these confidence scores, and the top *k* candidates (*k =* 1 or 5) are selected for evaluation.

GlycoBART was built upon the BARTForConditionalGeneration architecture from the BART model [37, 38]. BART is a sequence-to-sequence model that combines the advantages of BERT’s bidirectional context encoding with autoregressive decoding, making it effective for generative tasks. GlycoBART features a 12-layer encoder and a 12-layer decoder, each utilizing multi-head self-attention with 16 attention heads. The encoder operates similarly to that in GlycoBERT with 768-dimensional latent representations. The decoder then utilizes the resulting latent representations to autoregressively generate the corresponding Glycan sentence. During inference, beam search is used to generate multiple candidate glycan structures in parallel (n_beams_ = 32), from which the top *k* predictions are selected for evaluation.

GlycoBERT model contains 96 million parameters, while GlycoBART has 207 million. We trained two versions of each model. The first version was trained using an 85:15 train:test splits of the dataset to evaluate inference accuracy. For this version, spectra from the same MS/MS run could appear in both the training and test sets. The second version, denoted with -F suffix (GlycoBERT-F and GlycoBART-F), was trained using the entire dataset. All models were trained on four NVIDIA RTX A5000 GPUs using AdamW optimizer AdamW [39] optimizer with a learning rate of 1 × 10−^5^, with CrossEntropy loss function . GlycoBERT was trained for 50 epochs with a batch size of 256 while GlycoBART was trained for 30 epochs with a batch size of 80.

### Performance evaluation

We evaluated model performance on the test and independent datasets using four distinct, hierarchical levels of accuracy: mass, composition, topology, and structure.

1. *Mass Accuracy*: A prediction was considered correct if the absolute error between the monoisotopic masses of the predicted matches the annotated glycan mass with error less than 0.001 Da.
2. *Composition Accuracy*: This required the predicted glycan to have the same monosaccharide composition as the experimental annotation (*i*.*e*., the same count of each type of monosaccharide).
3. *Topological Accuracy*: A prediction was deemed correct if it shared the same monosaccharides and branching patterns as the experimental annotation, without considering glycosidic linkage information.
4. *Structural Accuracy*: (the most stringent metric) a prediction was structurally accurate if it matched the experimental annotation in both topology and glycosidic linkage.

For each of these four levels, the overall accuracy was reported as the percentage of predictions that satisfied the corresponding criterion. GlycoBERT’s accuracy was computed based on its top-1 prediction, i.e. the prediction receiving the highest score. In contrast, GlycoBART’s accuracy was evaluated for both top-1 and top-5 predictions. A top-*k* prediction was considered correct if the experiment annotation structure was present among the *k* predictions with the highest confidence scores.

### CANX KO Dataset for Independent Validation

HEK293T human embryonic kidney cells that underwent CANX knockout were obtained from previous work [40]. Cells were lysed in 1X invitrosol (Thermo Fisher) and 100 mM NH_4_HCO_3_, supplemented with 20% ACN. Lysis was followed by sonication at 62% amplitude with 10s pulse and 10s pause cycles, repeated three times with intermittent cooling on ice.

Proteins were reduced with 5 mM tris(2-carboxyethyl) phosphine at 37 °C for 45 min and subsequently alkylated with 10 mM iodoacetamide for 45 min at room temperature in the dark. Sixty-four (64) μg of the resulting proteins were immobilized on polyvinylidene difluoride (PVDF). N-glycans were then released via overnight incubation with 2U PNGase F (Promega, WI, USA) at 37 °C.

Released N-glycans were reduced with 1 M NaBH4 in 50 mM KOH for 3 h at 50 °C, then neutralized with equimolar acetic acid. Desalting and enrichment were performed using strong cation exchange resin Dowex 50WX8 (200-400 mesh) followed by PGC solid phase extraction.

Mass spectrometry analysis was carried out using post-column make-up flow PGC-LC-MS/MS platform in negative ion mode [41]. PGC-LC separation was performed using HyperCarb column (3 μm particle size, 180 μm ID x 100 mm length, Thermo Scientific) with a gradient of 10 mM NH_4_HCO_3_ containing either 0% (Solvent B) or 70% (v/v) acetonitrile (Solvent C) [42]. The gradient program was as follows: 5 % to 50% B from 0.001 to 0.1 minute, 50% to 0% B in next 0.9 minute, 0% to 14% B in next 0.1 minute, 14% to 26% B in next 48.9 minutes, 26% to 0% B in next 15 minutes, 0% to 100% B in next 4 minutes, held at 100% for next 9 minutes, 100% to 1% B in next 1 minutes, and finally held at 1% B for next 11 minutes. CID energy used was 33%.

We used GlycoBERT-F and GlycoBART-F to generate top-5 candidate glycan predictions. Subsequently, these predictions underwent a post-processing workflow to filter for correctness. The post-processing included the following steps:

a. *Mass check*: Glycan prediction monoisotopic mass was within 10 ppm of the experimental value.
b. *Glycan type check*: For this specific use case, only N-linked glycans belonging to the class Mammalia were considered valid for prediction.
c. *Diagnostic ion verification: For glycan predictions containing specific monosaccharides, including* Neu5Ac, Fuc, and GlcNAc, the corresponding diagnostic ions must be present among the top 20 most intense peaks in the mass spectrum.
d. *Fragment ions peaks match*: At least 3 of the top 15 de-isotoped peaks in the experimental spectrum must correspond to a theoretical fragment ion of the glycan prediction.

### Database Search using GlycoPAT

We adopted a variant of GlycoPAT (GlycoProteomics Analysis Toolbox) [43], where the mass calculations have been modified for glycans rather than glycopeptides. This variant is called Glycomics Analysis Toolbox (GlycomicsAT). Glycan annotations of MS/MS spectra were performed using this tool for the CANX KO dataset. Here, we first generated a custom library of N-glycans, encompassing bi-, tri-, and tetra-antennary structures, including those with core fucosylation and bisecting GlcNAc relevant to the CANX KO study. Based on this library, GlycomicsAT generated their theoretical fragment ions and experimental annotations.

For each experimental MS/MS spectrum, GlycomicsAT first identifies potential glycan-spectrum matches (GSM) by comparing the precursor m/z to the glycan library within a user-defined mass tolerance (tolerance: 10pmm). It then assigns an “Ensemble Score” (ES) to the candidate GSM using an algorithm that incorporates cross-correlation analysis between the experimental and theoretical spectra, the percentage of matched fragment ions (tolerance: 0.5Da), the number of the 10 most intense experimental peaks that align with theoretical ions, and Poisson probability-based p-value to assess statistical significance. To reduce false positives, GlycomicsAT employs a decoy-based strategy, generating scrambled monosaccharide masses, while keeping total glycan mass the same, to estimate false discovery rate (FDR). In our analysis, glycan-spectrum matches with an ensemble score greater than 0.4 and FDR<1% were preserved. Finally, the glycan annotations were manually inspected for the presence of diagnostic B/Y fragment ions in the MS/MS spectrum, and these manually validated GSMs were retained.

### Statistical analysis

Statistical analyses were conducted using SciPy (v.1.15.3) on Python 3.11.12.

## RESULTS

### Transformer Models for Glycan Structure Prediction from Tandem Mass Spectra

Mass spectrometry (MS) data encode information on glycan structures through the mass-to-charge (m/z) ratios and peak intensities of fragment ions. However, the interpretation of MS/MS spectra is inherently contextual: the meaning of any single peak depends on the presence and intensity of other peaks, as well as on the specific settings used during MS analysis. This interdependency among m/z peaks parallels the way context shapes meaning in natural language, thereby motivating the use of the transformer architecture— originally popularized by large language models like GPT—to MS data analysis.

To enable the application of transformer architectures for MS/MS glycan annotation, we transformed MS/MS spectra and glycan structures into “MS sentences” and “Glycan sentences”, respectively (see Methods). Each MS sentence is a sequence of words encoding information from MS analysis settings, normalized retention time, precursor mass, and MS peaks derived from a given spectrum (see Fig. 1A). We employed a binning strategy to convert continuous variables—including normalized retention time (RT) and m/z ratios—into discrete words. To incorporate information from MS peak intensities, the m/z peaks in each MS sentence are ordered in descending intensities. In this manner, peak intensities are captured by the positional embeddings in the transformer models (see Fig. 1C). On the other hand, a Glycan sentence represents a glycan structure as a series of its constituent antennae, separated by a whitespace (see Fig. 1B). Each antenna is encoded as a sequence of words, beginning with the terminal monosaccharide and followed by alternating linkage-monosaccharide pairs, terminating at the core monosaccharide (see Methods).

We explored two Natural Language Processing (NLP) tasks: sequence classification and conditional sequence generation, to predict glycan structures from MS/MS spectra (see Fig. 1C). For the sequence classification, we adapted the BERT architecture [36], specifically BertForSequenceClassifica2on, to create GlycoBERT, a multi-class classification model for predicting glycan structure from MS sentence. GlycoBERT’s bidirectional processing of input sequences is particularly suitable for capturing long-range dependencies within mass spectra, enabling the model to discern relationships between peaks even when they are distantly located in the sequence. For the conditional sequence generation approach, we adapted the BART architecture, specifically using BartForCondi2onalGenera2on, to build GlycoBART. The BART architecture is an extension of BERT that incorporates a decoder to endow generative capability. Conditional generation is commonly used in NLP for tasks such as text summarization, where the model generates a concise summary of a longer text. GlycoBART uses an MS sentence as the input sequence and outputs a Glycan sentence. For both GlycoBERT and GlycoBART, we developed custom vocabularies and tokenizer to handle the tokenization of MS and Glycan sentences.

Figure 1C illustrates the overall architecture and workflow of GlycoBERT and GlycoBART. An MS sentence, once tokenized, is converted into word and positional embedding forms for input to the encoder. In GlycoBERT, the encoder output is then passed to a classification head that assigns the spectrum to the most probable glycan class, finally outputting a ranked list of candidate glycans. In GlycoBART, the encoder output, together with the decoder input, serves as input to the BART decoder, which has been trained to generate glycan structure (sentence) predictions in an autoregressive manner. GlycoBART employs a beam search to generate several possible glycan sentences in parallel and identify the most likely glycan structure. Lastly, a post-processing step transforms the output Glycan sentence(s) into IUPAC format.

### Performance evaluation and comparison

We first assessed the accuracy of GlycoBERT and GlycoBART using a held-out test dataset. Figure 2A presents a comparison of mass, compositional, topological, and structural accuracies of glycan predictions generated by GlycoBERT, GlycoBART, and CandyCrunch [32]. Across all accuracy metrics, GlycoBERT consistently outperforms both GlycoBART and CandyCrunch. The top-1 glycan predictions from GlycoBART exhibited test accuracies comparable to those of CandyCrunch. However, when assessing the top-5 glycan predictions, GlycoBART demonstrated superior accuracies than CandyCrunch.

**Fig. 2.**
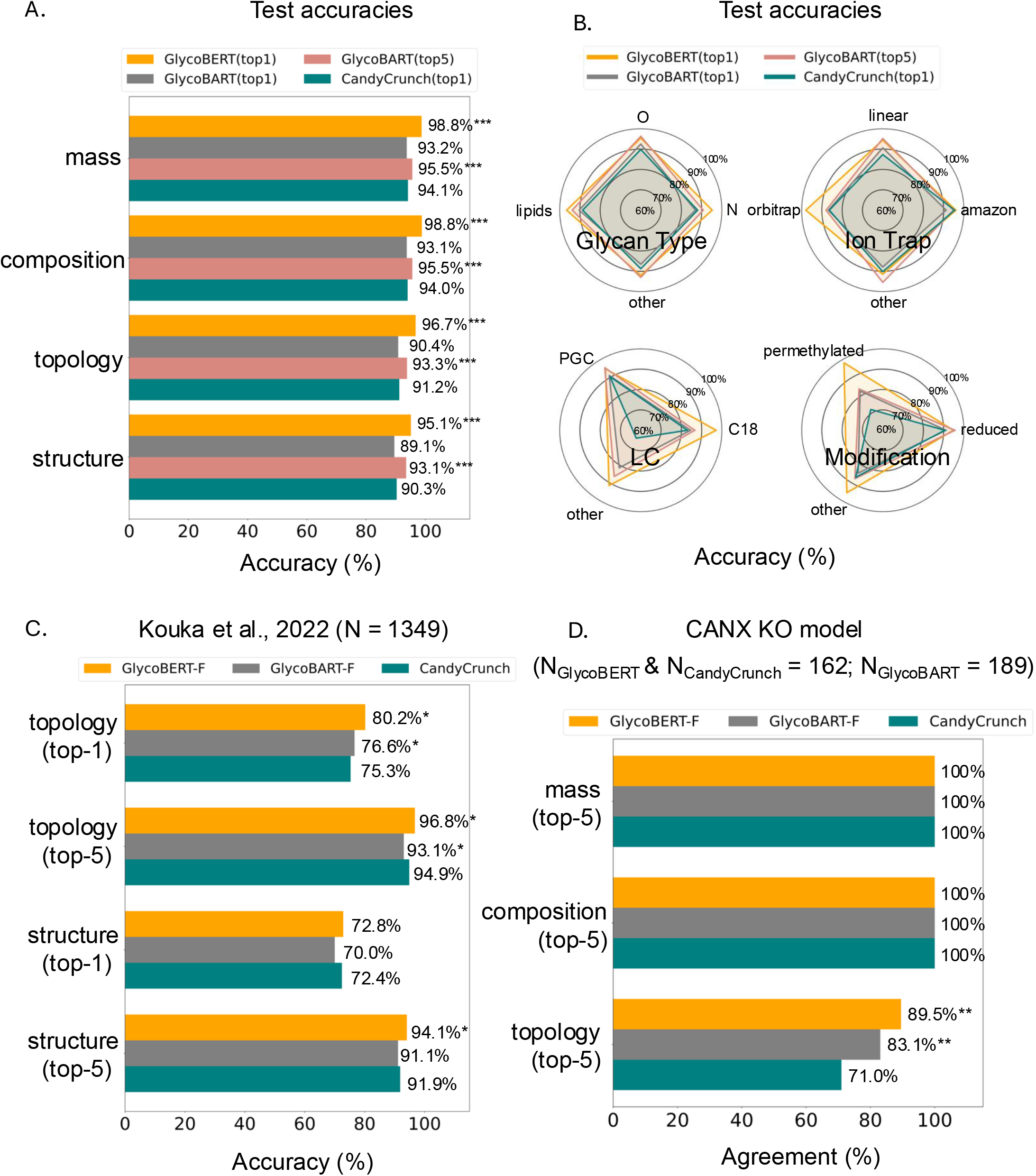
Performance evaluation of glycan structure prediction models. **(A)** Accuracy comparison on the held-out test dataset. Mass, compositional, topological, and structural accuracies are shown for predictions from GlycoBERT (top-1), GlycoBART (top-1 and top-5), and CandyCrunch (top-1). *** indicates *p* < 10−^5^, two-sided two-proportion z-test, comparing GlycoBERT or GlycoBART to CandyCrunch. **(B)** Structural accuracy stratified by experimental metadata. Structural accuracies from the test set are broken down by liquid chromatography (LC) method, glycan modification, ion trap, and glycan type. The ‘other’ category includes subtypes not present in the CandyCrunch model. **(C)** Validation on an independent external dataset. Topological and structural accuracies were evaluated on the Kouka *et al*. O-glycan dataset (*n* = 1,349 spectra), which was not used for model training or testing. * indicates p < 0.05, two-sided two-proportion z-test, comparing GlycoBERT-F or GlycoBART-F to CandyCrunch. **(D)** Agreement with database search method. Predictions from the deep learning models were compared against annotations from GlycoPAT on an N-glycan library from CANX-knockout HEK293T cells. Bars show the percentage agreement in mass, composition, and topology for spectra identified by both methods. The number of overlapping spectra is indicated for each comparison. ** indicates p < 0.01, two-sided z-test for proportions, comparing GlycoBERT-F or GlycoBART-F to CandyCrunch.

To gain deeper insights into model performance, we analyzed the test accuracies stratified by experimental and biological metadata, including liquid chromatography (LC) method, ion trap, glycan modification, and glycan type. Subtypes that are not present in the CandyCrunch model were grouped under the “other” category within each metadata group. As depicted in Fig. 2B, GlycoBERT and GlycoBART demonstrated more robust and consistent performance across various categories when compared with CandyCrunch. This was particularly evident for MS analyses using permethylated modification and LC methods besides PGC and C18 (*i*.*e*., ‘other’ category).

To further validate model performance beyond the test dataset, we compared GlycoBERT-F and GlycoBART-F—the version of these models trained using the full MS/MS dataset—with CandyCrunch on independent datasets from the Kouka et al. study [44]. This dataset consists of O-glycan profiles from 25 different CHO-K1 cell lines that were transiently transfected with various combinations of glycosyltransferases along with PSGL-1. It is important to note that MS/MS spectra from this external dataset were not part of the curated MS dataset used in the training of GlycoBERT-F and GlycoBART-F (see Methods). As shown in Fig. 2C, GlycoBERT achieved the highest topological and structural accuracies overall, outperforming GlycoBART and CandyCrunch. GlycoBART exhibited the lowest topological and structural accuracies with the exception of topological accuracy for top-1 predictions.

### Comparison to Database Search Method

We compared glycan predictions from deep learning (DL) based methods, GlycoBERT-F, GlycoBART-F, and CandyCrunch, with glycan annotations from a Glycomics data processing version of GlycoPAT [43] called GlycomicsAT, a database search method (see Methods for details). Specifically, we employed each of these methods to annotate MS/MS dataset from HEK293T kidney cell line that underwent CANX gene knock out (KO) from prior work [40]. The CANX gene encodes the calnexin chaperone protein, which is involved in the folding and quality control of N-linked glycoproteins within the endoplasmic reticulum [45]. The gene KO study was conducted to understand the role of Calnexin protein in N-linked glycosylation.

Agreement was quantified as a percentage of these spectra where at least one glycan prediction from the DL method matched a GlycomicsAT prediction in terms of mass, composition, or topology. Since GlycomicsAT predictions did not specify glycosidic linkages, structural comparisons were not performed. Fig. 2D indicates moderate to good agreement between GlycomicsAT and these DL-based methods. Notably, glycan annotations from GlycomicsAT showed better agreement with those generated by GlycoBERT-F and GlycoBART-F than with CandyCrunch (see Supplemental Data for full results).

### *De novo* glycan predictions by GlycoBART

A key advantage of GlycoBART among the other DL methods is its ability to perform *de novo* predictions—that is, to generate glycan structures not present in its training data. When applied to the CANX KO dataset, GlycoBART-F identified two such *de novo* structures.

The first *de novo* glycan structure, Neu5Ac(a2-6)Gal(b1-4)GlcNAc(b1-2)Man(a1-3)[Man(a1-2)Man(a1-3)Man(a1-6)]Man(b1-4)GlcNAc(b1-4)GlcNAc, corresponds to the composition h6n3s1. GlycoBART assigned this structure to a precursor ion with a mass of 1890.664 Da (m/z 945.3332, z = 2). This assignment is supported by B/Y fragment ions in the spectrum (see Fig. 3A), including B1 (s1, m/z 290.13), Y9 (h6n3, m/z 1600.68), B3 (h1n1s1, m/z 655.09), and Y7 (h5n2, m/z 1235.65).

**Fig. 3.**
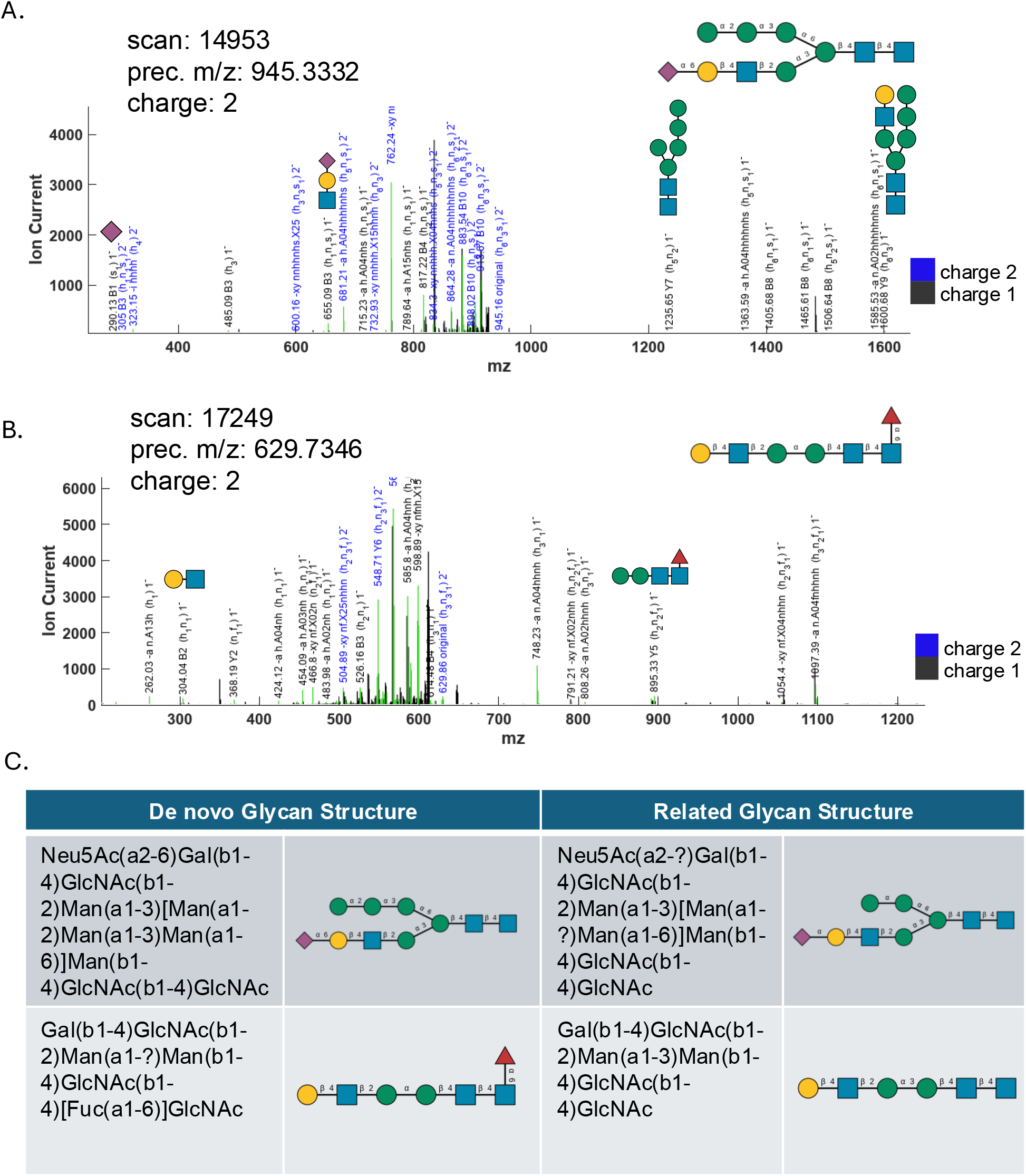
Validation of *de novo* glycan structures predicted by GlycoBART. **(A)** MS/MS spectrum of a precursor ion (m/z 945.3332, z=2) associated with the de novo glycan h6n3s1. Key fragment ions supporting the assignment are annotated. **(B)** MS/MS spectrum of a precursor ion (m/z 629.7346, z=2) associated with the de novo glycan h3n3f1, with supporting fragment ions annotated. **(C)** The de novo structures (left) are shown with structurally related glycans (right) that were also identified in the CANX KO dataset. The presence of these structural neighbors, differing by a monosaccharide residue, support the *de novo* assignments.

The second *de novo* structure, Gal(b1-4)GlcNAc(b1-2)Man(a1-?)Man(b1-4)GlcNAc(b1-4)[Fuc(a1-6)]GlcNAc, has the composition h3n3f1, assigned a precursor ion with a mass of m/z 1259.4692Da (m/z 629.7346, z=2). This identification was supported by the B2 (h1n1, m/z 304.04) and Y5 (h2n2f1, m/z 895.33) fragment ions (see Fig. 3B).

To corroborate the above de novo inference, we searched the remaining GlycoBART glycan annotations for this MS/MS dataset for structurally related glycans. Our search revealed neighbors for both *de novo* structures (see Fig. 3C). For the first glycan (h6n3s1), we identified a variant with an additional mannose residue. For the second (h3n3f1), we found a corresponding structure that lacked a core fucose. The detection of these closely related glycans lends additional support to the de novo identifications.

To determine if these *de novo* predictions represented previously documented glycan structures, we performed structure-based searches in the GlyCosmos [47], GlyConnect [48], and UniCarbDB [49] databases. We found a match for the first structure Neu5Ac(a2-6)Gal(b1-4)GlcNAc(b1-2)Man(a1-3)[Man(a1-2)Man(a1-3)Man(a1-6)]Man(b1-4)GlcNAc(b1-4)GlcNAc in GlyCosmos (GlyTouCan ID: G36721QZ), indicating prior evidence. In contrast, the second glycan structure yielded no matches, suggesting a possible novel glycan or partially degraded N-glycan existing in the CANX cells.

## DISCUSSION

The accurate and efficient determination of glycan structures from tandem mass spectra remains a central challenge in glycomics, owing to the vast diversity, isomeric complexity of glycans and absence of gold standard data. In this study, we present GlycoBERT and GlycoBART, transformer-based models that bring a new way to enhance annotation accuracy and also generative capability for *de novo* glycan structure inference from MS/MS data. Central to our methodology is a novel representation of spectra and glycan structures as tokenized sequences, enabling the models to learn intricate relationships among spectral peaks, their intensities, and contextual experimental parameters. Unlike CNN-based approaches such as CandyCrunch that discretize the m/z range into bins and aggregate peaks within each bin [32], our MS sentence representation preserves individual peak information through positional encoding. The transformer architecture’s self-attention mechanism is particularly effective at capturing complex, long-range dependencies in spectral data. Our transformer-based models, in particular GlycoBERT, outperform CandyCrunch across different accuracy metrics and demonstrate robust performance across diverse experimental parameters. The consistent performance across different glycan types (N-, O-, lipid-linked, and free glycans) and experimental conditions suggests that transformer-based approaches can provide a robust framework for glycomics analysis. This is particularly important given the field’s reliance on diverse analytical platforms and sample preparation methods, which have historically made cross-study comparisons challenging.

A key advantage of GlycoBART is its generative capability, which enables the prediction of glycan structures not present in the training set. This capability addresses a fundamental limitation of classification-based methods that are inherently constrained to previously known structural classes. This generative approach is particularly valuable for glycomics, where the theoretical diversity of possible structures far exceeds what has been experimentally characterized and catalogued in databases. Our analysis of the CANX knockout dataset illustrates this potential: GlycoBART not only matched known structures but also proposed two de novo glycans, one of which was absent from major glycan databases, potentially representing a novel structure. The ability to discover such structures has important implications for understanding disease mechanisms, as aberrant glycosylation patterns are hallmarks of various pathological conditions [9, 10, 11].

While GlycoBERT and GlycoBART represent significant advances in glycan structure prediction, several challenges and opportunities remain. The quality and consistency of training data, as highlighted in the CandyCrunch study [32], directly impact model performance. Future efforts should focus on curating larger, more diverse, and better-annotated datasets. While transformer models benefit from large-scale pretraining, our current dataset size limited us to training models for specific tasks: sequence classification and conditional sequence generation. Pretraining on larger, unlabeled spectral datasets, as is common in NLP and proteomics, could further enhance model performance and generalizability, but would require access to much larger and more diverse MS/MS datasets.

Looking ahead, exploring more powerful architectures, such as hybrid models combining transformers with graph neural networks—which could explicitly represent glycan structures—or models incorporating domain-specific knowledge (e.g., biosynthetic rules), may yield further gains in predictive performance. With access to larger unlabeled datasets, pretraining strategies such as masked language modeling could be employed to learn richer representations of spectral data, which can then be fine-tuned for glycan prediction tasks. Developing methods to interpret model predictions—such as attention visualization and feature attribution—will be important for building user trust and facilitating biological discovery. Finally, integrating these computational advances with experimental glycomics workflows, including real-time prediction during data acquisition and automated validation protocols, represents an important step toward making transformer-based glycan analysis a routine component of glycomics research.

## CONCLUSIONS

In this study, we addressed the challenge of accurate and high-throughput glycan structure determination from tandem mass spectrometry data. By conceptualizing MS/MS spectra and glycan structures as tokenized sequences, we successfully developed transformer-based models, GlycoBERT and GlycoBART, to tackle this complex problem. Our results demonstrate that this approach yields significant improvements in performance. GlycoBERT achieves state-of-the-art accuracy in classifying known glycan structures, outperforming existing CNN-based method CandyCrunch. More significantly, the generative capability of GlycoBART overcomes the limitation of classification models by enabling *de novo* inference of glycan structures. The successful identification of two novel structures in an independent dataset, one of which a previously unreported glycan, highlights the potential of such generative model for novel discovery. Overall, the development of GlycoBERT and GlycoBART represents a step forward for computational glycomics. These models provide a robust and generalizable framework that is less constrained by experimental parameters and is not limited to a predefined library of known structures.

## ACKNOWLEDGEMENT

The authors gratefully acknowledge funding support from the National Heart, Lung, and Blood Institute (grant number HL103411). They also appreciate the valuable discussions with Drs. Rebekah Gundry and Wenjuan Zha regarding mass spectrometry analysis.

## DATA AND MODEL AVAILABILITY

Tandem mass spectrometry (MS/MS) dataset and glycan sentences (i.e. sequences of glycan antennae) are available on Zenodo (https://doi.org/10.5281/zenodo.15741423). The Python codes for training and glycan inference are available from a GitHub repository called glycoTrans (https://github.com/cabsel/glycotrans). The full version of GlycoBERT and GlycoBART models are available on HuggingFace (https://huggingface.co/CABSEL).

